# Bivalent chromatin as a therapeutic target in cancer: An *in silico* predictive approach for combining epigenetic drugs

**DOI:** 10.1101/2020.10.15.340679

**Authors:** Tomás Alarcón, Josep Sardanyés, Antoni Guillamon, Javier A. Menendez

## Abstract

Tumour cell heterogeneity is a major barrier for efficient design of targeted anti-cancer therapies. A diverse distribution of phenotypically distinct tumour-cell subpopulations prior to drug treatment predisposes to non-uniform responses, leading to the elimination of sensitive cancer cells whilst leaving resistant subpopulations unharmed. Few strategies have been proposed for quantifying the variability associated to individual cancer-cell heterogeneity and minimizing its undesirable impact on clinical outcomes. Here, we report a computational approach that allows the rational design of combinatorial therapies involving epigenetic drugs against chromatin modifiers. We have formulated a stochastic model of a bivalent transcription factor that allows us to characterise three different qualitative behaviours, namely: bistable, high- and low-gene expression. Comparison between analytical results and experimental data determined that the so-called bistable and high-gene expression behaviours can be identified with undifferentiated and differentiated cell types, respectively. Since undifferentiated cells with an aberrant self-renewing potential might exhibit a cancer/metastasis-initiating phenotype, we analysed the efficiency of combining epigenetic drugs against the background of heterogeneity within the bistable sub-ensemble. Whereas single-targeted approaches mostly failed to circumvent the therapeutic problems represented by tumour heterogeneity, combinatorial strategies fared much better. Specifically, the more successful combinations were predicted to involve modulators of the histone H3K4 and H3K27 demethylases KDM5 and KDM6A/UTX. Those strategies involving the H3K4 and H3K27 methyltransferases MLL2 and EZH2, however, were predicted to be less effective. Our theoretical framework provides a coherent basis for the development of an in-silico platform capable of identifying the epigenetic drugs combinations best-suited to therapeutically manage non-uniform responses of heterogenous cancer cell populations.

**Author summary:** Heterogeneity in cancer cell populations is one of the main engines of resistance to targeted therapies, as it induces nonuniform responses within the population that clears the sensitive subpopulation, whilst leaving unaffected the non-responsive cells. Although this is a well-known fact, few successful approaches have been proposed aimed at both quantifying the variability associated to cell heterogeneity, and characterising strategies that circumvent its drug-resistance inducing effects. Here we present a computational approach that addresses these issues in the particular context of targeting epigenetic regulators (specifically, chromatin modifiers), which have been proposed as therapeutic targets in several types of cancer and also in ageing-related diseases. Our model predicts that the more successful combinations involve modulators of demethylase activity (specifically, KDM5/6 and UTX). By contrast, strategies involving EZH2 activity are predicted to be less effective. Our results support the use of our framework as a platform for *in silico* drug trials, as it accounts for non-homogeneous response of cell populations to drugs as well as identifying which subpopulations are more likely to respond to specific strategies.

## Introduction

Heterogeneity is a primary cause of *de novo* and acquired resistance to targeted treatments in cancer therapeutics [1–3]. There is ample evidence of genetic diversity among cancer cells both within and across tumours have been shown to exist [4–6], with genomic instability being the main engine of genetic heterogeneity [7]. However, it is now apparent that genetic variability alone does not account for the whole variation in responses to targeted anti-cancer drugs [8, 9]. Non-genetic heterogeneity has two broad sources, epigenetic and stochastic, the latter being due to intrinsic factors such as noise in gene expression [10–13] and asymmetric cell division [14, 15], and it can produce phenotype diversity even in genetically identical cells [8, 9, 16]. Not surprisingly, the analysis of the consequences of non-genetic cell-to-cell variability in the cellular response to drugs and its potential impact for the treatment of human diseases including cancer has become a crucial issue to understanding drug resistance phenomena and developing new targeted agents [1, 3, 17].

Beyond its well-documented role in development [18–20], bivalent chromatin is an emerging epigenetic trait of cancer that has been identified as a druggable therapeutic target [19, 21]. Genomic studies have found that there exists significant overlap between bivalent domains in pluripotent stem cells and regions in cancer exhibiting abnormal patterns of methylation, which pinpoints bivalency as a promising marker of tumourigenesis. Furthermore, different bivalent chromatin regulators have been identified as targets for cancer therapeutics [22–24]. An outstanding example of such regulator is EZH2 (a histone methyl transferase which is the catalytically active component of the PRC2 complex), which has found been to be mutated or overexpressed in a number of cancers [23]. Specifically, somatic EZH2 mutations have a major role in promoting lymphoid transformation in germinal-center leukemias [21]. Another family of epigenetic regulators that has emerged as a potential target are the so-called lysine demethylases (KDMs), whose expression has been found to be dysregulated in several types of neoplasms [22, 24]. Flavahan et al. [25] have recently proposed bivalent chromatin regulators as key factors in epigenetic plasticity, i.e. the epigenetically-driven alteration of the stability of cellular states (phenotypes). Variations in the activity of certain epigenetic enzymes such as EZH2 can lead to *repressive* states, where the epigenetic barriers are raised resulting in cells locked in a certain state, regardless of the presence of signalling cues instructing them to exit such states. By contrast, such alterations can also produce *permissive* states in which the epigenetic barriers are lowered so that cellular states change by the mere presence of random noise. The latter states often lead to pathological plasticity and cell reprogramming [25–27].

The so-called Waddington or epigenetic landscape, a conceptual framework allowing the integration of both genetic and non-genetic variability, was proposed over seven decades ago [28]. In analogy to the potential energy landscape of physical systems, Waddington put forward a representation of a cellular state in the form of an *effective potential landscape* where the local minima correspond to cellular phenotypes. Phenotypic switch transitions occur when cells transits to other minima by jumping over the barriers separating adjacent basins. Such transitions are driven by intrinsic or extrinsic fluctuations [29, 30]. Noteworthy, the topography of such landscapes depends on a complex set of biochemical interactions and, in particular, on the values of the corresponding set of rate constants [17, 31–34]. These constants strongly depend on protein structure (e.g. the accessibility of a binding domain), which is encoded within the DNA [32]. Thus, an epigenetic landscape emerges from a particular genotype [17], and therefore can be interpreted as a representation of the genotype-phenotype map [35–40]. Beyond their dependence on DNA sequence, in the case of biochemical reactions involving enzymatic catalysis, the associated rate constants depend also on the concentration of metabolic cofactors [33, 34], which may differ among cells thus adding a further source of cell-to-cell heterogeneity.

In order to shed some light on the role of bivalent chromatin as a therapeutic target, we propose a mathematical model of a simple gene regulatory circuit with a bivalent promoter (see Figures 1(a) and (b)). Specifically, our model addresses mutual nonlinear feedbacks between marked nucleosomes and transcription factors (TFs), which drive not only the rates of both addition of new marks and TF synthesis [41–45], but also the heterogeneity of cellular populations [17, 33, 34, 46, 47].

**Figure 1.**
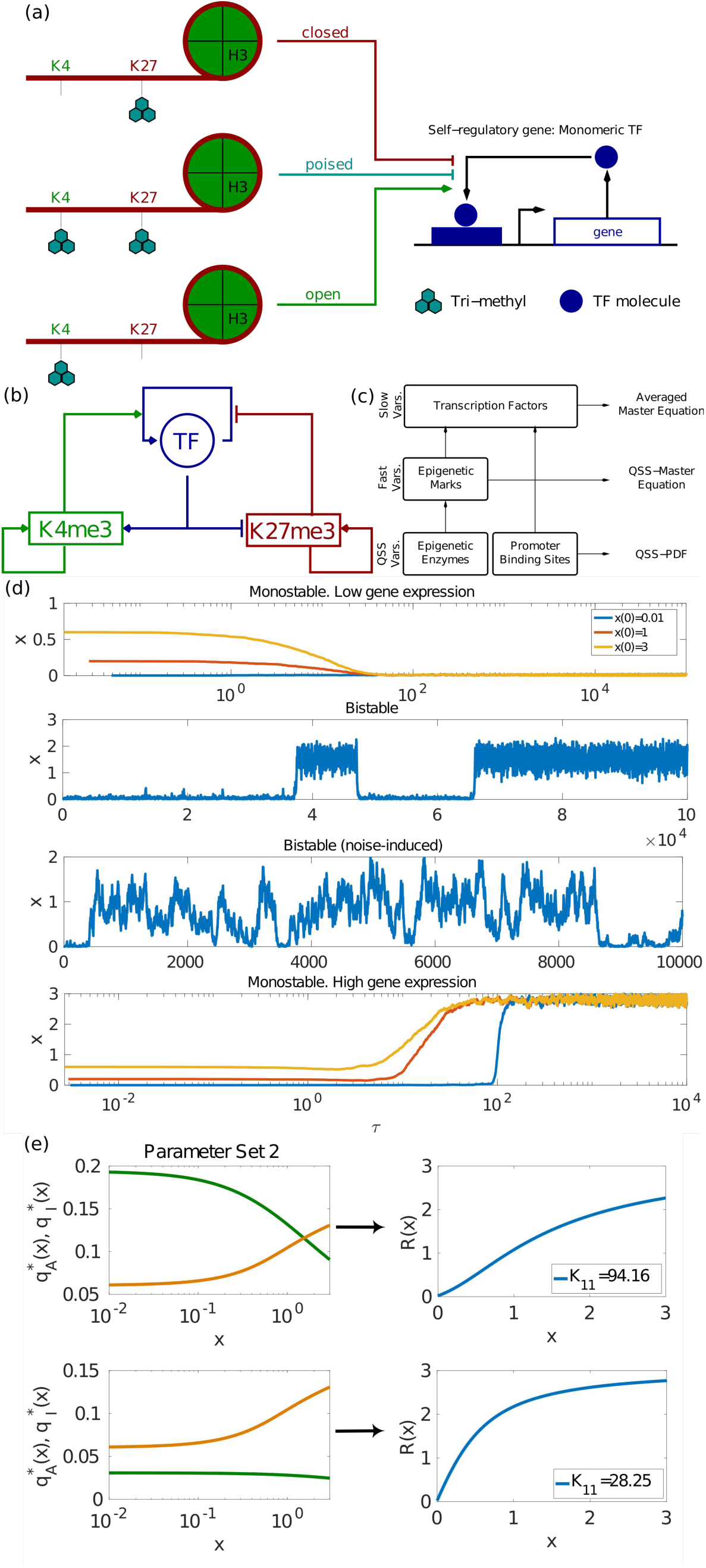
Summary of the model formulation and model results. Panel (a) shows a schematic representation of the epigenetic-regulatory modifications considered in our model (i.e. trimethylation of H3K4 and H3K27), which affect promoter/enhancer regions. We focus on a transcription factor (TF) which is self-regulatory and monomeric. This panel also shows the different states of bivalent chromatin (*open* (*closed*) if H3K4me3 (H3K27me3) predominates, *poised* if both marks are present) and how they affect TF expression. Plot (b) depicts a scheme showing the feedbacks between ER modifications and TF (full details given in Section S.1.1, S1 File). Plot (c) displays a schematic representation of the model reduction approximation, based on the presence of multiple time scales. Such a separation of time scales allows for a hierarchical elimination of the faster variables (full details given in Section S.1.2, S1 File). Plot (d) shows results of direct simulation of the process (carried out by means of the Gillespie algorithm [48, 49]). These simulations illustrate the four type of behaviours afforded by the stochastic model: in descending order, inactive TF, bistability, noise-induced bistability, and active TF. Finally, Plot (e) illustrates one of our main results, namely, that the TF exhibits emergent, non-linear behaviour (e.g. bistability) which could not be observed in the absence of bivalent chromatin. Specifically, we show that such behaviour is associated to the tunability of the effective cooperativity, *n*_*eff*_, of the TF regulatory circuit: by altering the activity of the chromatin-modifying enzymes we can control the quantity *n*_*eff*_. In the particular case shown in the figure, by varying *K*_11_ (which corresponds to the *K*_*M*_ of the KDM that removes H3K27me3), the effective cooperativity changes from *n*_*eff*_ > 1 (upper plot) to *n*_*eff*_ ≤ 1 (lower plot). The response curve, *R*(*x*), and the *n*_*eff*_ of the TF are defined by 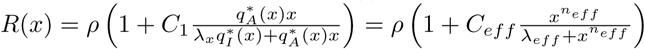.

The determination of epigenetic regulatory mechanisms has triggered an interest in developing mathematical models regarding both epigenetic regulation (ER) of gene expression [27, 33, 34, 41–43, 45, 50–53] and epigenetic memory [50–52, 54–58]. Specifically, Sneppen et al. [41] were the first to propose a computational model to address the coupling between transcription factor and epigenetic regulation and how it produces ultrasensitive behaviour, but they did not consider bivalent chromatin. Such a coupling has been further studied in the context of bivalent chromatin [42, 43, 45]. Although these works, similarly to the model presented here, describe bistability in gene regulatory systems with TF self-regulation, it should be noted that they consider a self-activating TF with high intrinsic cooperativity (*n* = 3) [42, 43]. Moreover, noise-induced phenomena in gene regulatory networks have been extensively studied [12, 59–63]. In particular, Biancalani and Assaf [12] have studied noise-induced bistability in a regulatory network (toggle switch) of monomeric, self-activating, mutually-inhibiting genes, but they did not consider ER.

Our model (whose main features are summarised in Figure 1) proves that bivalent chromatin regulation can induce complex behaviour even in the simplest circuits of gene regulation, which would not exhibit such features in the absence of bivalent regulation. We consider a model of a self-regulatory, monomeric TF with a bivalent promoter (see Figure 1(a)), characterised by a number of both positive and negative feedbacks identified in [42, 43] (and schematically illustrated in Figure 1(b)). The resulting model, in spite its apparent simplicity, exhibits considerable complexity regarding its nonlinearities and multiplicity of time scales. In order to make analytical progress, we exploit the latter and, by means of an asymptotic model reduction and singular perturbation analysis, we reduce the model to a one variable stochastic system (Figure 1(c)). By doing so, we have been able to determine, both numerically and analitycally, that the system exhibits three different behaviours: monostability, bistability, and noise-induced bistability [64] (Figure 1(d)). Specifically, we show that the activity of the epigenetic enzymes directly controls the effective cooperativity of the gene regulatory circuit, which impinges on its ability to produce bistable behaviour (Figure 1(e)).

Our analysis of a bivalent TF is based on the explicit calculation of the Waddington landscape for the system under consideration, which allows us to exhaustively classify the set of possible landscape topographies. We account for the effects of heterogeneous behaviour in the epigenetic-regulatory system [46, 47] by means of an ensemble approach previously developed [33, 34], in which we take advantage of the direct connection between landscape topography and the rate constants that characterise the underlying biochemical networks from which it emerges. After validating our ensemble approach against quantitative experimental data [20, 21, 65], we finally address the issue of what epigenetic strategies are more effective regarding their ability to overcome heterogeneity. We conclude that those involving tackling the activity of demethylases are more efficient than those that targeting the activity of methyltransferases.

## Materials and methods

### Model formulation

Here, we focus on an epigenetic regulation (ER) model that consists of four enzymatic reactions, which account for the addition and removal of a 3-methyl, to be referred as “me3”, epigenetic mark to two specific lysine residues in the tail of the H3 histone (H3K4 and H3K27, see Figure 1(a)). Each of these enzymatic reactions is modelled by means of the well-known Michaelis-Menten (MM) model of enzymatic catalysis (see [66] and S1 File for details specific to our model). We apply this model to address how specific enzymes catalyse the addition of the me3 mark to each residue (MLL2 to H3K4 and EZH2 to H3K27 [25]), and the removal of me3 from modified residues (specifically, KDM2/KMD5 from H3K4me3 and KDM6 from H3K27me3 [67, 68]). Beyond the regular mechanisms of enzyme kinetics, epigenetic regulation exhibits several feedback mechanisms that need to be taken into account [41, 54]. Modified residues enhance recruitment of epigenetic enzymes, providing a positive feedback mechanism where residue modification increases the rate of further me3 addition (see Figure 1(b)). Following previous models of ER [33, 34], we account for such positive feedbacks by taking the rate constants associated to each of the me3-addition MM reactions to change linearly with the number of corresponding modified residues (see Table S.1 in S1 File). Furthermore, recruitment of epigenetic enzymes to H3K4 is also enhanced by the presence of transcription factors [41–43] (see Figure 1(b)). Such positive feedback is also accounted for by assuming the rate constants of the MM reaction associated with H3K4→H3K4me3 to change linearly with the number of TF molecules. By contrast, TF presence hinders recruitment of me3-addition to H3K27 [42, 43] (see Figure 1(b)). To take into account this negative feedback, we have assumed that the rate corresponding to the back reaction of the MM reaction associated to H3K27→H3K27me3 depends linearly on the number of TF molecules (see Table S.1 in S1 File).

Regarding the gene regulatory system, we consider a simple self-regulatory gene (see Figure 1(a)), i.e. a gene such that its protein stimulates its own expression, modelled by a simple Hill model (with Hill constant *α* = 1, since we are considering a monomeric TF) [69] with natural decay and basal production of the TF molecule. As shown in Figure 1, the feedback between TF dynamics and ER is accounted for by assuming that the rate of TF binding to the gene’s promoter region is proportional to the number of H3K4me3, whilst the rate TF unbinding is proportional to the number of H3K27me3. A fully detailed account of the model formulation is provided in Section S.1 in S1 File.

### Asymptotic analysis and model reduction

The model resulting from the previous discussion is rather complicated. In order to make analytical progress, we exploit the presence of separation of time scales. In particular, we resort to asymptotic model reduction techniques and singular perturbation analysis (see Figure 1(c) for a schematic representation). A fully detailed derivation is presented in Section S.1 in S1 File, which we summarise here. We proceed with our analysis in two steps. First, by considering the usual assumptions regarding separation of time scales and the associated quasi-steady state approximations (QSS approximation) normally involved in the analysis of MM and Hill systems (see [66, 69], and S1 File, Section S.3, for the stochastic version of the QSS approximation), one can considerably simplify the system.Specifically, under assumptions considered in detail in S1 File, we can reduce the dynamics of the whole system to that of three variables, namely, the number of TF molecules and the number of H3K4me3 and H3K27me3 residues. In the case of the mean-field limit, it reduces to the following system of three differential equations (see Section S.1 in S1 File for a detailed derivation):

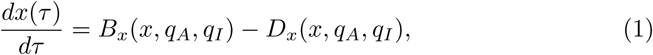

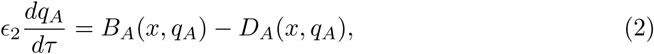

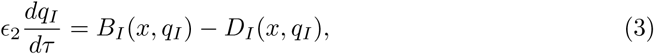

where

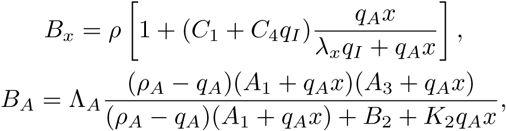

and

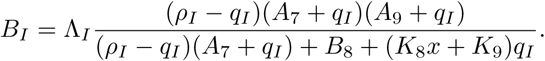

Also,

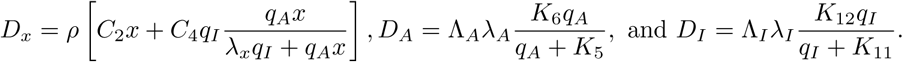

The quasi steady-state (QSS) concentration of H3K4me3, 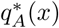; and of H3K27me3, 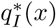, are the solutions of 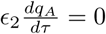 and 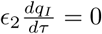, respectively.

Note that, stemming from the assumption that 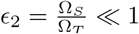, where Ω_*S*_ and Ω_*T*_ are characteristic scales for the number of H3K4 and H3K27, and TF molecules, respectively, the resulting reduced system, Eqs. (1)–(3), still exhibits slow-fast features (see Section S.1 in S1 File). The same holds true for the stochastic version of the model, which is described by the corresponding Master Equation (ME):

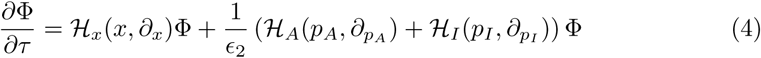

where the operators 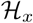, 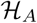, and 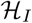 are given by

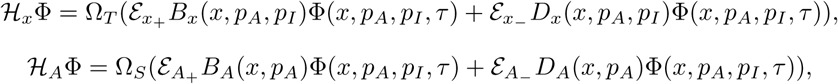

and

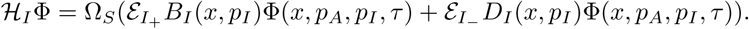

The step operators are defined in the usual manner: 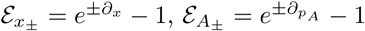, and 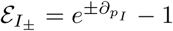 [70,71].

In order to proceed forward we propose that the PDF Φ(*x, p*_*A*_, *p*_*I*_, *τ*) be expanded in powers of 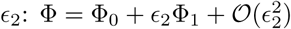 [64, 72]. In S1 File, we show that, at the lowest order, Φ_0_ can be written as Φ_0_(*x, p*_*A*_, *p*_*I*_, *τ*) = *ϕ*_*x*_(*x,τ*)*ϕ*_*A*_(*p*_*A*_|*x*)*ϕ*_*I*_(*p*_*I*_|*x*), with *ϕ*_*A*_ and *ϕ*_*I*_ such that 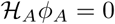 and 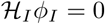. The WKB solution to the latter equations is given by

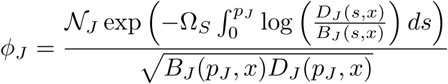

with *J* = *A, I*. However, as usual with singularly-perturbed problems such as Eq. (4), the lowest order approximation does not provide any information regarding *ϕ*_*x*_(*x, τ*) and we need to resort to the next order [72, 73]. At steady state, by considering the appropriate solvability conditions, we can show that, to the lowest order, 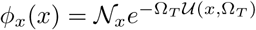, where the effective potential, *𝓤*, is given by:

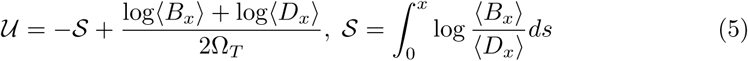

where 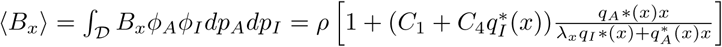 and 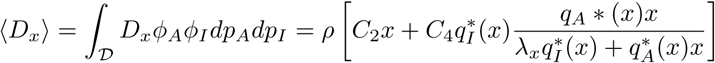.

A fully detailed derivation of the asymptotic calculation is given in S1 File, Section S.1.2.

### The ensemble approach: Modelling heterogeneity and parameter sensitivity analysis

Our ensemble approach serves two distinct but not unrelated purposes, namely, to perform a parameter sensitivity analysis and to model heterogeneity observed in epigenetic regulatory systems [46, 47]. Regarding the former, given the complexity of our model, we need to consider carefully robustness to variations in parameter values. We consider such issue using an ensemble approach introduced in [33, 34]. Briefly, we generate an ensemble of parameter sets. A parameter set is described by means of a vector, *θ*, defined as (see Table S.2 in S1 File):

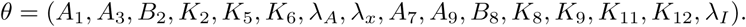

Each component of *θ* is independently generated by sampling from a uniform prior distribution within a certain prescribed range. Randomly generated parameter sets are accepted only if the *q*_*A*_ and *q*_*I*_ are monostable (see Section S.1.4 in S1 File). The sets of parameter values thus generated are then classified into several sub-ensembles depending on the number of roots of 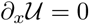, i.e. the number of modes of *ϕ*_*x*_, and 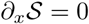, i.e. the number of mean-field steady-states. We consider a monostable sub-ensemble composed by those *θ*s such that both 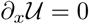 and 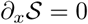 have only one real, positive root. We split this set of parameters further by considering a high gene expression (high GE) sub-ensemble if the modal point *x*_*_ ≥ *x*_*T H R*_; and a low gene expression (low GE) sub-ensemble *x∗ < xTHR*. We consider a bistable sub-ensemble composed by those *θ*s such that both equations have three roots. Last, the noise-induced bistable sub-ensemble comprises *θ*s sych that 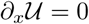 has three roots while 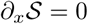 has only one root. Representative examples of landscapes associated with systems, i.e. *θ*, belonging to each of four sub-ensembles are shown in Figure 2. We then analyse the ensembles in terms of the comparison of the marginal posterior PDFs for each of the components of *θ* and also their correlations. We expect that by discerning which parameters present statistically significant differences between sub-ensembles, we can identify which of them are key to produce the behaviour associated to a particular sub-ensemble [33, 34]. We refer the reader to S1 File, Section S.1.4.2, for a full account of the details of the classification procedure and further statistical analysis.

**Figure 2.**
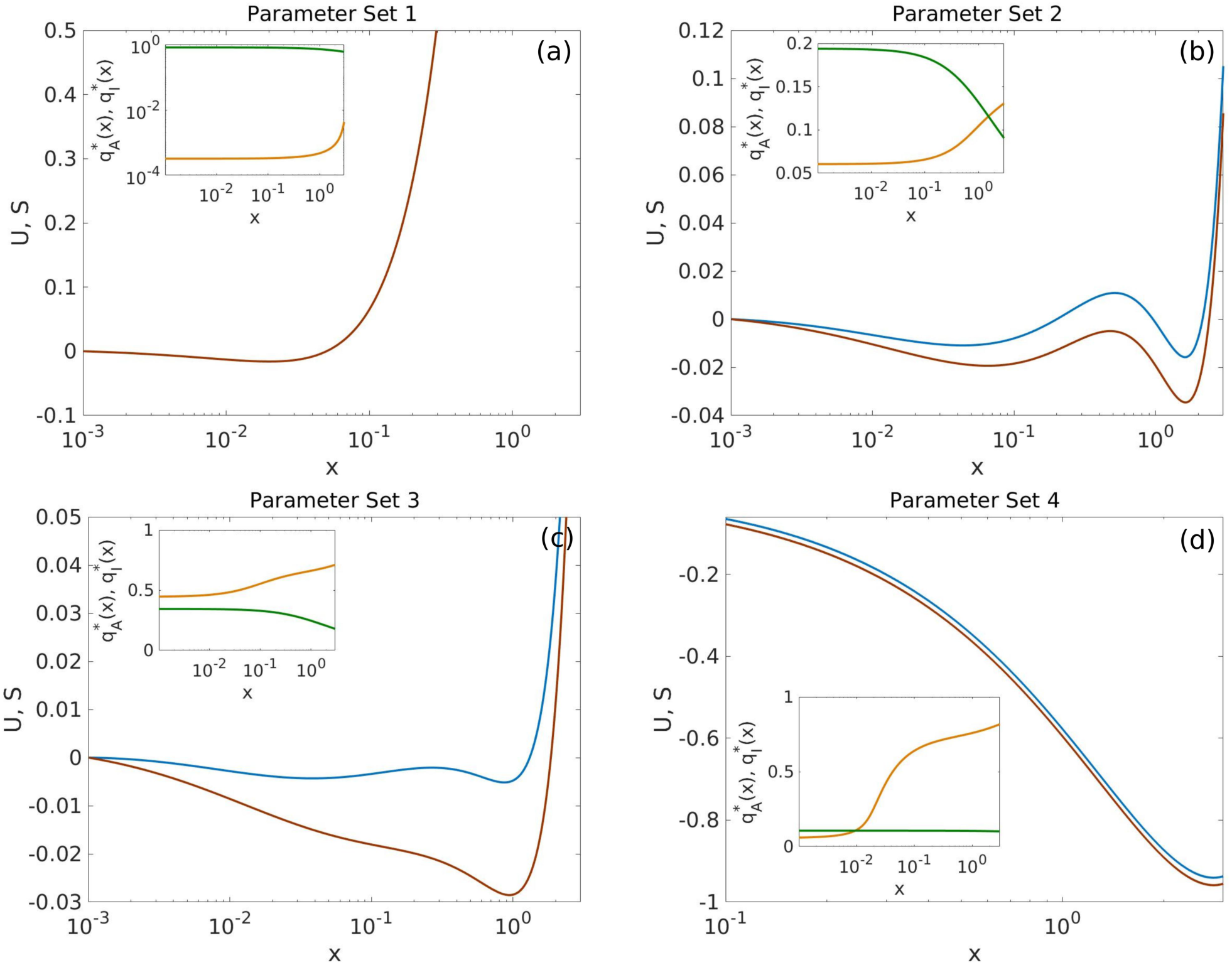
Representative examples of the four sub-ensembles. Plot(a) shows an example of a monostable, low gene expression system, belonging to the Low GE sub-ensemble. Plots (b) and (c) show examples of systems within the bistable and the noise-induced bistable sub-ensembles, respectively. Plot (d) shows an example of a monostable, high gene expression system, belonging to the high GE sub-ensemble. See main text for details. Colour code: Red lines show 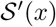, blue lines correspond to *𝓤′*. Green and orange lines within the insets represent 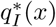 and 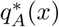, respectively. Parameter values are given in Tables S.3-2.6 of S1 File.

Experimental evidence of heterogeneity in ER systems comes from a number of sources. It has been directly observed in single-cell CHIP-seq experiments [47]. Other studies have provided evidence that the *de novo* reprogramming potential is higher within selected subpopulations of cells and that such pre-existing epigenetic heterogeneity can be tuned to make cells more responsive to reprogramming stimuli [46]. Furthermore, Beguelin et al. [21] have reported that the efficiency of drugs that target chromatin regulators, specifically EZH2 inhibitors, is very heterogeneous: their IC_50_ varies from one cell type to another in two orders of magnitude (see figure S.1 in the supplementary information of Ref. [21]). In order to mimic, at least in part, the existing ER heterogeneity within a cell population from a particular tissue, the generated ensemble of ER systems can be used to identify properties at the population level. Our approach follows closely that of Ref. [33]. The above procedure provides us with an ensemble of parameter sets, *θ*, that are compatible with each of the four behaviours we have found (Low GE, High GE, bistability, and noise-induced bistability). In order to check whether the heterogeneity within the ensemble is compatible with existing biological variability, we compare with quantitative experimental data provided by Beguelin et al. [21].

## Results

### Validation of the ensemble model

We first validate our approach against qualitative and quantitative results [20, 21, 25, 65, 74]. Figures 3(a), (b), and (c) show summary statistics of the monostable and bistable sub-ensembles. Specifically, Figure 3(a) shows the distribution of modes of *ϕ*_*x*_(*x*), *x*_∗_, within the monostable subsensemble. As we can see this distribution is bimodal, which supports the definition of two monostable sub-ensembles: the low gene expression (Low GE) sub-ensemble and the high gene expression (High GE) one.

**Figure 3.**
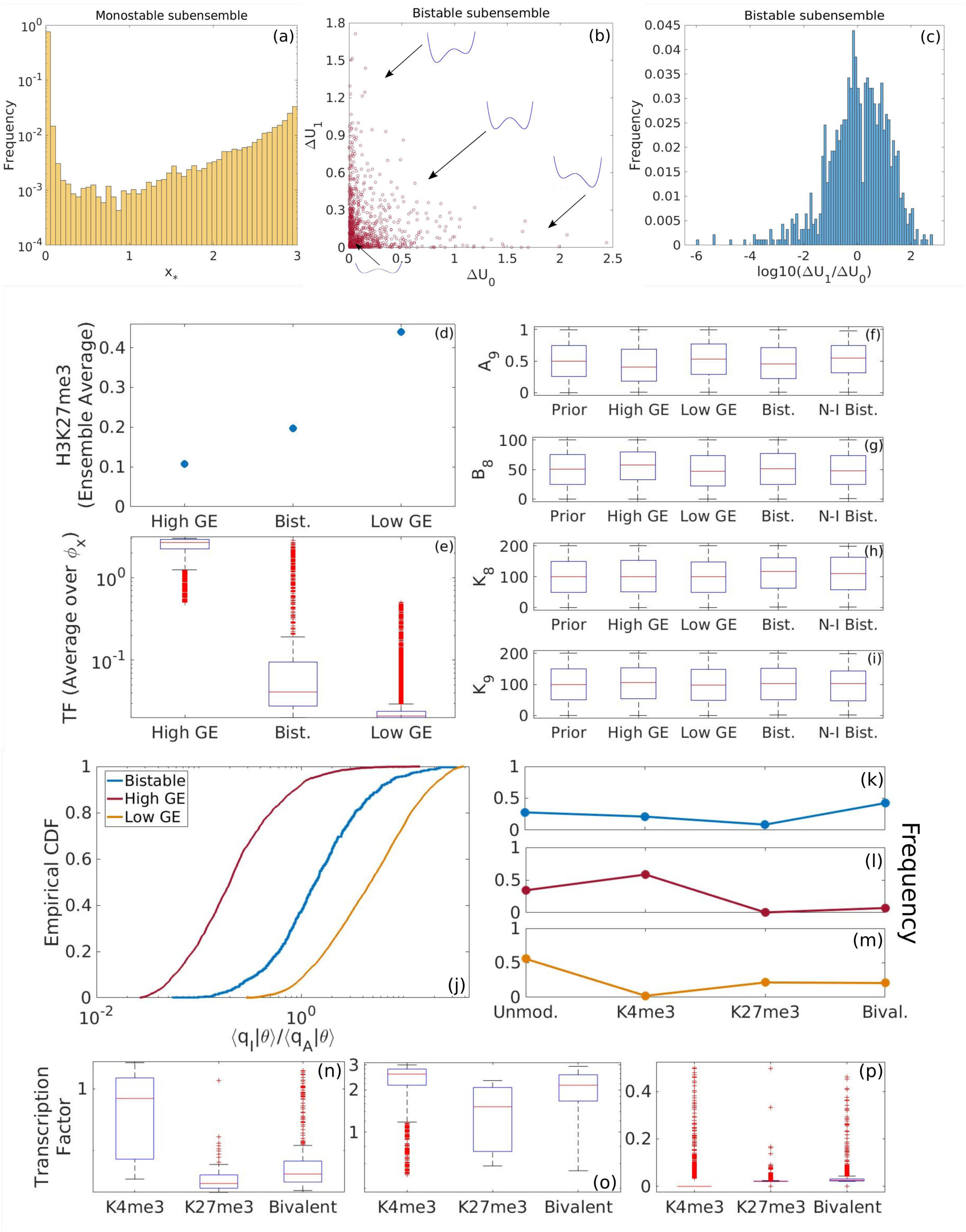
Validation of the ensemble model. Plots (a), (b), and (c) show summary statistics regarding the monostable and bistable sub-ensembles. Plot (a) shows the distribution of modes, *x*_∗_, within the monostable sub-ensemble. Plot (b) shows the scatter plot of the effective potential barriers, ∆*𝓤*_0_ and ∆*𝓤*_1_, within the bistable sub-ensemble. Plot (c) shows the histogram of the ratio ∆*𝓤*_1_/∆*𝓤*_0_ within the bistable sub-ensemble. Panels (d) and (e) show data regarding statistics of the Bistable, High GE, and Low GE sub-ensembles. Plot (d) displays sub-ensemble averages of the H3K27me3 marks and Plot (e) shows the boxplots of the TF levels (specifically, we use the modal TF concentration, *x*_∗_ as a proxy) for each of the sub-ensembles. These data are to be compared with the experimental results from [21]. Plots (f)-(i) show the boxplots for the distributions of kinetic parameters of EZH2 for each of the sub-ensembles. Finally, plots (j) to (p) report statistics of the Bistable, High GE, and Low GE sub-ensembles regarding their comparison to quantitative data provided in [20]. Plot (j) show the empirical cumulative density function (CDF) for the ratio 〈*q*_*I*_|*θ*〉_*x*_/〈*q*_*A*_|*θ*〉_*x*_ < 5) for each of the sub-ensembles. Plots (k), (l), and (m) show the proportion of elements of each sub-ensemble with no detectable methylation marks (which we define as those systems such that 〈*q*_*A*_|*θ*〉_*x*_ < 0.05 and 〈*q*_*I*_|*θ*〉_*x*_ < 0.05), dominant K4me3 marks 〈*q*_*I*_|*θ*〉_*x*_/〈*q*_*A*_|*θ*〉_*x*_ < 0.8), dominant H3K27me3 marks 〈*q*_*I*_|*θ*〉_*x*_/〈*q*_*A*_|*θ*〉_*x*_ < 5), and bivalent marks. These are to be compared with data from figure 2 in [20]. Plots (n), (o), and (p) show the modal TF expression, *x*_∗_, for H3K4me3 marked systems, H3K27me3 marked systems, and bivalent systems for the bistable, High GE, and Low GE sub-ensembles, respectively. These results are to be compared with figure 4 in [20].

Figure 3(b) shows a scatter plot of the effective potential barriers, ∆𝓤_0_ and ∆𝓤_1_, within the bistable sub-ensemble. According to [25], there are several pathological situations that can be described on the basis of such quantities. Repressive states, associated with higher barriers, and permissive states, corresponding to lower barriers, appear in response to dysregulation of chromatin modifiers. Both states are represented in our ensemble by those ER systems which lay at the tails of Figure 3(b) (or within the tails of the histogram shown in Figure 3(c)). Other anomalies of the epigenetic landscape involve lowering of both barriers which facilitate noise-driven exploration of different phenotypes. These systems are also represented in our bistable ensemble by those systems located in the lower left corner of Figure 3(b), i.e. such that ∆*𝓤*_0_ and ∆𝓤_1_ are both small and of the same order of magnitude.

Beyond qualitative arguments, quantitative validation is possible. Consider the data from Bernstein et al. [20] regarding the number and relative expression levels of genes with bivalent marks, H3K4me3-dominated genes, H3K27me3-dominated genes, and unmarked genes (methylation status). Such data was provided for both embryonic stem cells (ESCs) and several types of differentiated cells (DCs) (shown in Figures 2 and 4 in [20], respectively). Figures 3(j)–(m) address our results regarding methylation status patterns. Figures 3(k)–(m) show results of the frequency of each methylation status for all three sub-ensembles. The pattern exhibited by the Bistable sub-ensemble is the same as the ESCs data from Ref. [20] (see figure 2 in Ref. [20]): the frequency of unmarked, bivalent and H3K4me3-dominated is of the same order, whilst the frequency of negatively (H3K27me3) marked systems is much lower. High GE sub-ensemble shows the same pattern as DCs: the frequency of positive marks is much increased with respect to ESCs, whereas the frequency of bivalent marks is considerably reduced, to the same level as the frequency of negative marks. By contrast, the Low GE pattern is not compatible with any of the patterns reported by Bernstein et al.

**Figure 4.**
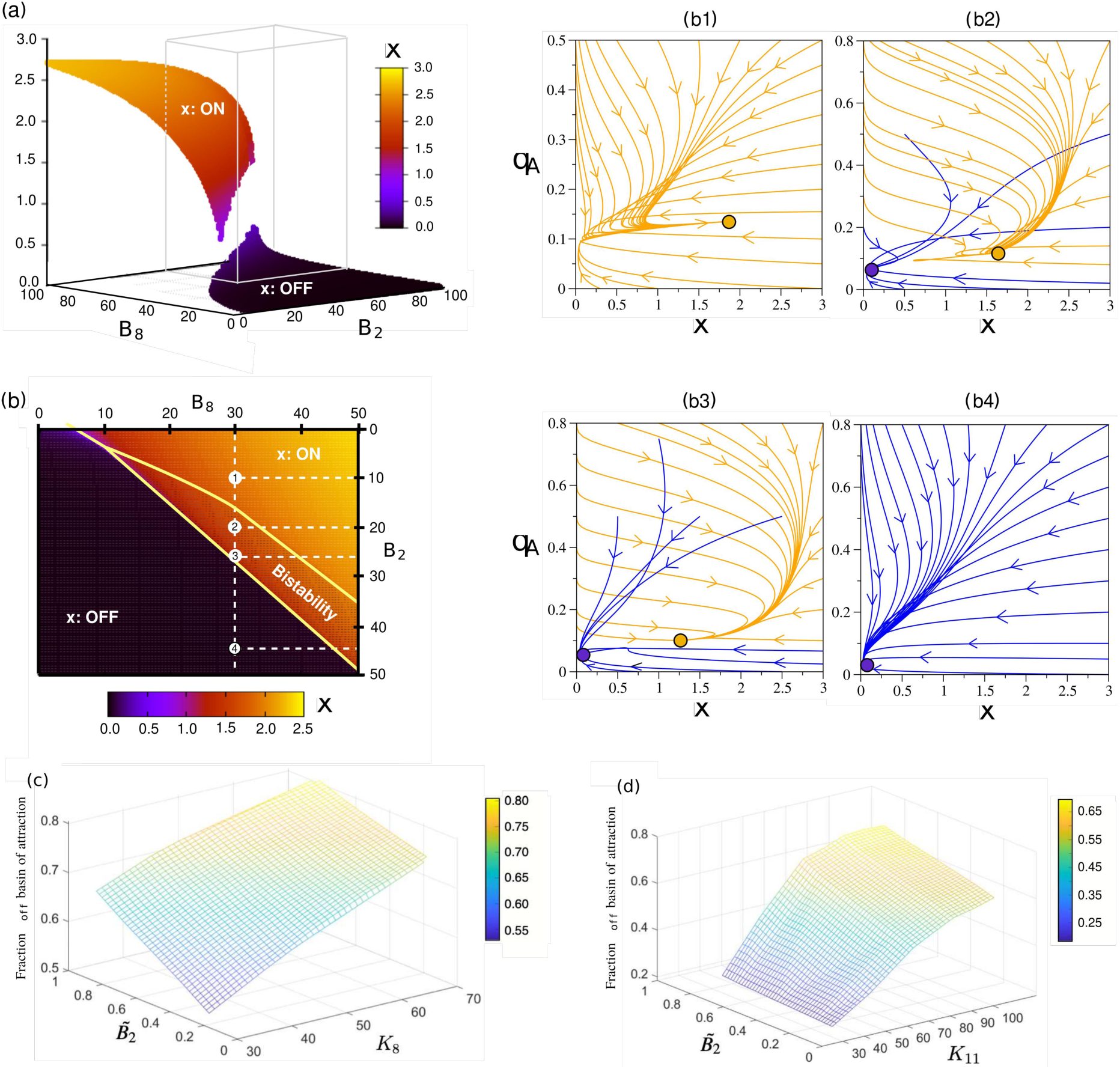
Equilibria in the parameter space (*B*_2_, *B*_8_). Panel (a) shows the equilibrium concentration of the TF (variable *x*) in the parameter space (*B*2, *B*8), the upper (resp., lower) sheet corresponding to the high (resp., low) level of expression, labelled as ON (resp., OFF) state. Panel (b) displays an enlarged view within the range contained in the cube in (a), namely 0 ≤ *B*_2_, *B*_8_ ≥ 50. The region contained in the central orangish area corresponds to bistability. (b1-b4) Phase portraits showing the dynamics in the parameter space (*x, q*_*A*_) for the combination of parameters *B*_2_, *B*_8_ indicated with dashed lines in (b), computed using *ϵ* = 0.1. The orange and blue orbits correspond to the dynamics achieving the ON and OFF states, respectively. Note that both (b2) and (b3) fall into the region of bistability. Panels (c) and (d) show the relative size of the basin of attraction of the OFF equilibrium for parameter sets within the bistability region: (c) Parameter plane 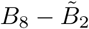, where *K*_11_ is set to be 94.155549; (d) Parameter plane 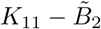, where *B*_8_ is set to be 30. For the sake of visualisation, 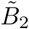 stands for a normalization of parameter *B*_2_, explained in Section S.2 in S1 File. Other constants are fixed according to the parameter set 2 (see Table S.7, Section S.2 in S1 File).

Figures 3(n)–(p) address the comparison with the levels of TF expression. We use the distribution of modal TF expression, *x∗*, for each methylation status. Figure 3(n) shows that the patterns of the Bistable sub-ensemble match the experimental behaviour of ESCs (see figure 4 in Ref. [20]), as the expression of H3K4me3-marked TFs is between 1 and 2 orders of magnitude larger than the levels exhibited by bivalent and H3K27me3-marked TFs. We also observe that TF expression is largely suppressed in bivalent systems, their levels being only slightly larger than that of H3K27me3-marked TFs. The pattern of TF expression within the High GE expression shown in Figure 3(o) fits that of DCs: H3K4me3-marked systems exhibit higher TF expression than that of H3K27me3-marked and bivalent systems, but the overall levels of expression are much more uniform than in the case of the Bistable sub-ensemble. The pattern of TF expression of the Low GE ensemble shown in Figure 3(p) are not consistent with any of the results of Bernstein et al.

Further validation of the ensemble model can be obtained by comparing to Beguelin et al. [21], who studied the role of somatic mutations of EZH2 on lymphoid transformation in germinal centre leukemias. They determined that EZH2 mutants, specifically Y641F and Y641N, lock B cells in an undifferentiated state during the germinal centre reaction. Such undifferentiated state can eventually lead to unchecked cell duplication (by silencing the cell-cycle inhibitor p27) and malignant progression. Beguelin et al. quantified the levels of H3K27me3 in cells carrying the wild-type (WT) EZH2 and cells carrying the mutants Y641F and Y641N (Figure 3b in [21]). They also collected quantitative information of the relative levels of expression of certain differentiation TFs (specifically IRF4, Figure S.4 in [21], supplementary information). These data can be compared with our ensemble model, as shown in Figures 3(d) and (e).

Figure 3(d) shows the sub-ensemble average of the number of H3K27me3-marked residues (see S1 File, Section S.1.5 for details) for the Bistable, High GE, and Low GE sub-ensembles. The predicted levels of H3K27me3 in the Bistable sub-ensemble doubles those in the High GE. This is consistent with the relative increase of H3K27me3 in the Y641F and Y641N mutants relative to WT (although in absolute numbers, our model prediction is slightly lower than those measured experimentally, figure 3b in [21]). The Low GE sub-ensemble exhibits a 4-fold increase in H3K27me3 with respect to the High GE sub-ensemble, which is out of the scale observed experimentally. Figure 3(e) shows boxplots of TF expression (specifically, of the modal level of expression, *x∗*) for each sub-ensemble. Comparing the results for the High GE and Bistable sub-ensembles, we notice that, consistently with Beguelin et al. (figure S.4 in the supplementary information of [21]), there is between 1 and 2 orders of magnitude of difference, average TF levels being higher in the High GE.

Finally, Figures 3(f)–(i) show boxplots for four parameters related to the enzymatic kinetics of EZH2 (see Eq. (3)) for each of the four sub-ensembles (see S1 File, Section S.1.4 for full analysis). By inspection, whilst there are obvious differences between the Low GE PDFs and those corresponding to the High GE and the Bistable sub-ensembles, the difference between the latter ones is much less obvious. In fact, quantitative analysis (see S1 File) shows statistically significant differences, but with *p*-values very close to the significance level. This observation is consistent with direct *in vitro* measurements of the Michaelis-Menten constant, *K*_*M*_, and the catalytic constant, *k*_*cat*_, of the wild-type EZH2 and Y641F and Y641N mutants by [65], who found that in all variants of the enzyme, the *K*_*M*_s retained similar values within the error bars.

Taken together, these results validate two essential aspects of our model. First, the ensemble variability appears to be consistent with the amount of heterogeneity observed in the experiments carried out in [20, 21, 65]. Furthermore, these results allow us to identify the systems within the Bistable sub-ensemble (and probably also the Noise-Induced Bistable one) with ESCs/undifferentiated cells, whereas the High GE systems can be identified with differentiated cell types. The Low GE sub-ensemble exhibits properties which are incompatible with either type.

### Bivalency allows for bistability and tunable cooperativity

Our analysis reveals that our model of a bivalent TF exhibits an array of complex behaviours, which are inaccessible to the gene regulatory system in the absence of bivalent ER (see [41]), and give rise to a variety of topographies of the epigenetic landscape, *𝓤*_*eff*_. Such behaviours are illustrated in Figures 1(d) and 2. In particular, besides the expected monostable behaviour (with both low and high TF expression, Figures 2(a) and (d), respectively), Figure 2(b) shows that the bivalent TF model exhibits bistability. Such behaviour is associated with effective cooperativity *n*_*eff*_ > 1 (see Figure 1(e)), in contrast to the monomeric self-regulating gene with no bivalent ER (which has *n*_*eff*_ = 1). By means of a sensitivity analysis, we determine that variation of key parameters associated with the activity of ER enzymes, specifically, *B*_2_, *B*_8_, *K*_5_ and *K*_11_ (see Eqs. (2)–(3)), provides us with ability to control *n*_*eff*_ thus allowing to suppress bistable behaviour, since variations of *n*_*eff*_ can drive the system through a saddle-node (s-n) bifurcation, thus removing pathological equilibria (see Figure 4). Such result is consistent with current experimental evidence, whereby controlling the activity of the ER enzymes, specifically of EZH2 [21, 75], has the effect of reducing pathologically-heightened epigenetic barriers and allow for physiological differentiation. Specifically, Figure 4 shows the change of the steady state concentration of TF as *B*_2_ and *B*_8_ vary. The s-n bifurcation causes sharp transitions at both the lower and upper yellow boundaries of Figure (4)(b), which enclose the region of bistability. The variety of dynamical behaviours exhibited by our system is depicted by several examples shown in the panels: (b1) monostability with TF at ON state (high level of expression); (b2)-(b3) with bistability behaviour; and (b4) monostability with the TF at OFF state (low level of expression).

In the case of bistable behaviour, it is interesting to analyse the variation of the basin of attraction (BA) of the high (ON) and low (OFF) TF concentration steady states as the activity of the chromatin-modifying enzymes changes. Specifically, for concreteness, we analyse how the BAs change as *B*_2_ and *B*_8_ vary (see Figure S.2 of S1 File), which impinge upon the activity of MLL2 and EZH2, respectively. Within the bistability region, the probability of reaching either state is sensitive to parameter variation. In Figure (4)(c) and (d), we show the results of exploring the region of bistability within the parametric space (*B*_2_, *B*_8_, *K*_11_) by computing the relative size, *P*_*off*_, of the basin of attraction of the OFF state (note that *P*_*off*_ + *P*_*on*_ = 1) using a Monte Carlo procedure. This quantity provides an approximation of the probability that a given combination of TF molecules and methylated histones evolve towards the OFF state. From Figure (4), we show that, as we increase the value of any of the three factors (*B*_2_, *B*_8_ and *K*_11_), the system is more likely to evolve towards an OFF state. This result illustrates that ON-OFF transitions can occur upon variation of the activity of epigenetic-regulatory enzymes in a deterministic way.

From the mechanistic point of view, a necessary (but not sufficient) condition for bistable behaviour (*n*_*eff*_ ≥ 1) is that the QSS mark concentrations, 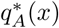 and 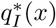 (Eqs. (2)–(3) and S1 File), exhibit the *X*-like pattern we observe in Figures 1(e) and 2(b). When this condition is satisfied, if *x* fluctuates below its crossover value, the system enters a regime that favours a continued decrease in gene expression since the negative marks dominate over the positive ones, and viceversa, thus inducing ultrasensitive behaviour associated with *n*_*eff*_ ≥ 1 and bistability. From this perspective, the mechanism behind the role of the activity of the ER enzymes is clear: by changing the value of key parameters, we can alter such pattern and induce transitions in the system (s-n bifurcations) from bistable to monostable behaviour (Figures 2(a) and (d)).

### Bivalency causes noise-induced bistability

Besides bistable behaviour, the bivalent TF model also exhibits noise-induced bistability, i.e. the ability of the system to sustain two stable steady-states only if the system is affected by noise. An example of this behaviour is shown in Figure 2(c), where we show that *ϕ*_*x*_(*x*) is bimodal for finite system size (Ω_*T*_ = 200). Such behaviour is lost for vanishing levels of noise in the system (when Ω_*T*_ → ∞). This feature is confirmed by direct Gillespie simulations of the stochastic process described by Eq. (4). Results are displayed in Figures 1(d) and 5, where we present simulations of the system for different system sizes, which show that, in agreement with our analytical approximation, noise-induced bistability ensues when the system size is reduced below its predicted critical value. Furthermore, we have ascertained that variations in ER enzyme activity induces noise-induced bistable behaviour. Figure S.5 in S1 File shows that modification of parameters controlling ER enzyme activity (specifically, *B*_2_, *B*_8_, and *K*_11_) remove mean-field bistability but preserve bimodality of the PDF (i.e. bistable behaviour is noise-induced). However, noise-induced bistability remains a scarcely robust phenomenon within the bivalent TF model: the percentage of noise-induced bistable *θ*s within the generated ensemble is ∼ 0.4% (see Table S.4 in S1 File) and the percentage of bistable *θ*s that respond to parameter change by becoming noise-induced bistable barely reaches a 10% of the size of the bistable sub-ensemble (see Figure S.5 in S1 File).

**Figure 5.**
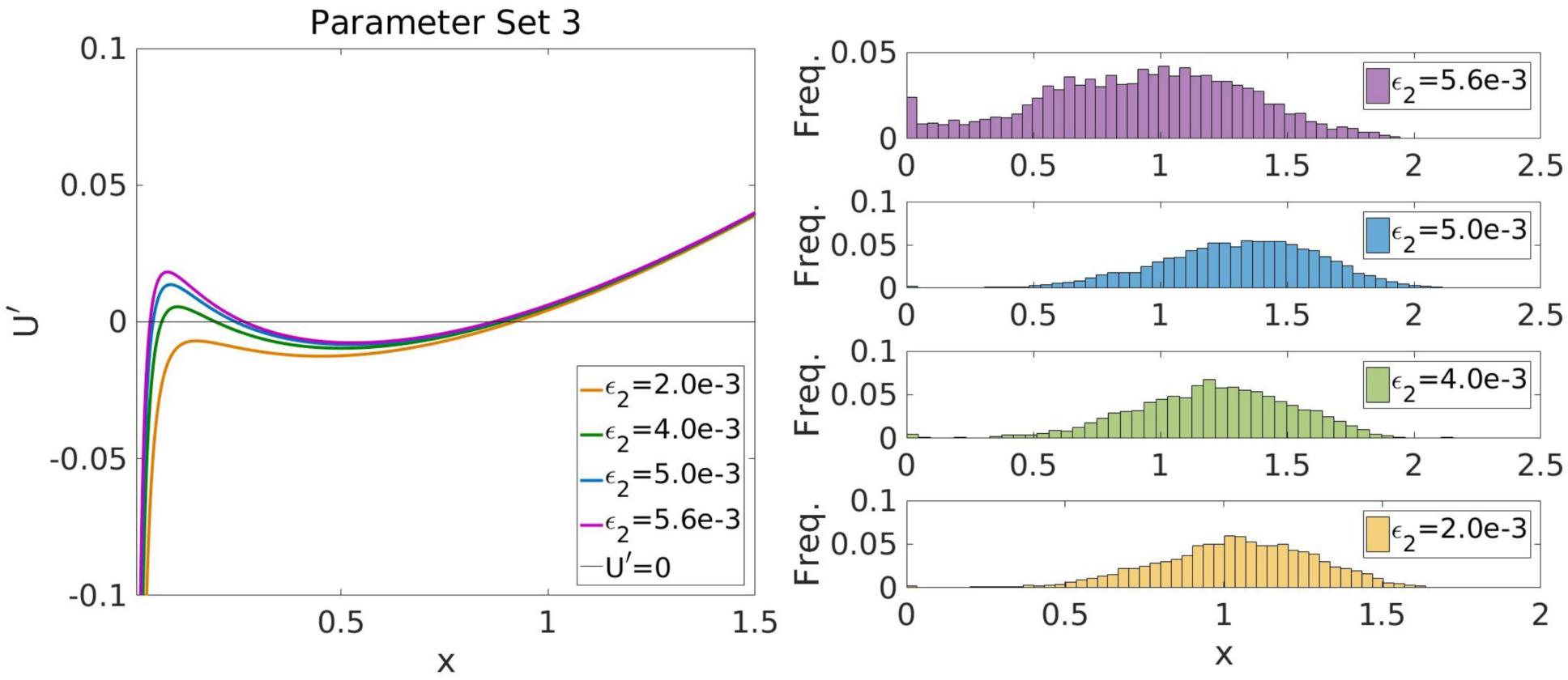
Noise-induced transition to bistable behaviour. This figures shows stochastic simulation results supproting the analytical prediction that our system exhibits a noise-induced transition form mono- to bi-stable behaviour. The plot shows how, as Ω decreases (i.e. the level of noise increases), the PDF of the system undergoes a transition from mono- to bi-modality.

### Tackling heterogeneity

Heterogeneity in cellular populations is one of the main obstacles regarding resistance to targeted therapies in cancer [1, 3]. Since the effects of drugs is not uniform on a heterogeneous cell population, those cells whose response is poorer become a resistant subpopulation that compromises drug efficiency. Heterogeneity has thus become a major factor in developing strategies to circumvent the emergence of resistance [76]. ER enzymes such as EZH2 have been recently proposed as targets to treat certain types of B cell leukemias [21, 75]. Since recent single-cell studies have revealed that heterogeneity also occurs in ER status [47], it is necessary to evaluate its effect on therapies that target ER enzymes.

Following [33, 34], where we put forward that heterogeneity in ER enzyme activity can be traced back to heterogeneity in the metabolic state of the cells, we analyse the effect of heterogeneity in terms of the response of the sub-ensemble of pathological *θ*-systems to changes in parameters that are relevant to ER enzyme activity. In this case, based on our previous discussion that allows us to identify systems within the Bistable sub-ensemble as compatible with the class of undifferentiated cells (see Figure 3(k)), we analyse the response of the Bistable sub-ensemble to three generic strategies: altering the activity of H3K4-modifying enzymes, altering the activity of the H3K27-modifying enzymes, and mixed strategies involving several combinations (see Figures 6(c), (d), and (e), respectively). The sensitivity of the sub-ensemble to changes in parameter values, or combinations therein, is measured as the proportion of systems that become monostable (i.e. the proportion of systems that change from the Bistable into to either the High GE or the Low GE systems). Our results are shown in Figure 6 (see also Section S.1.4 in S1 File).

**Figure 6.**
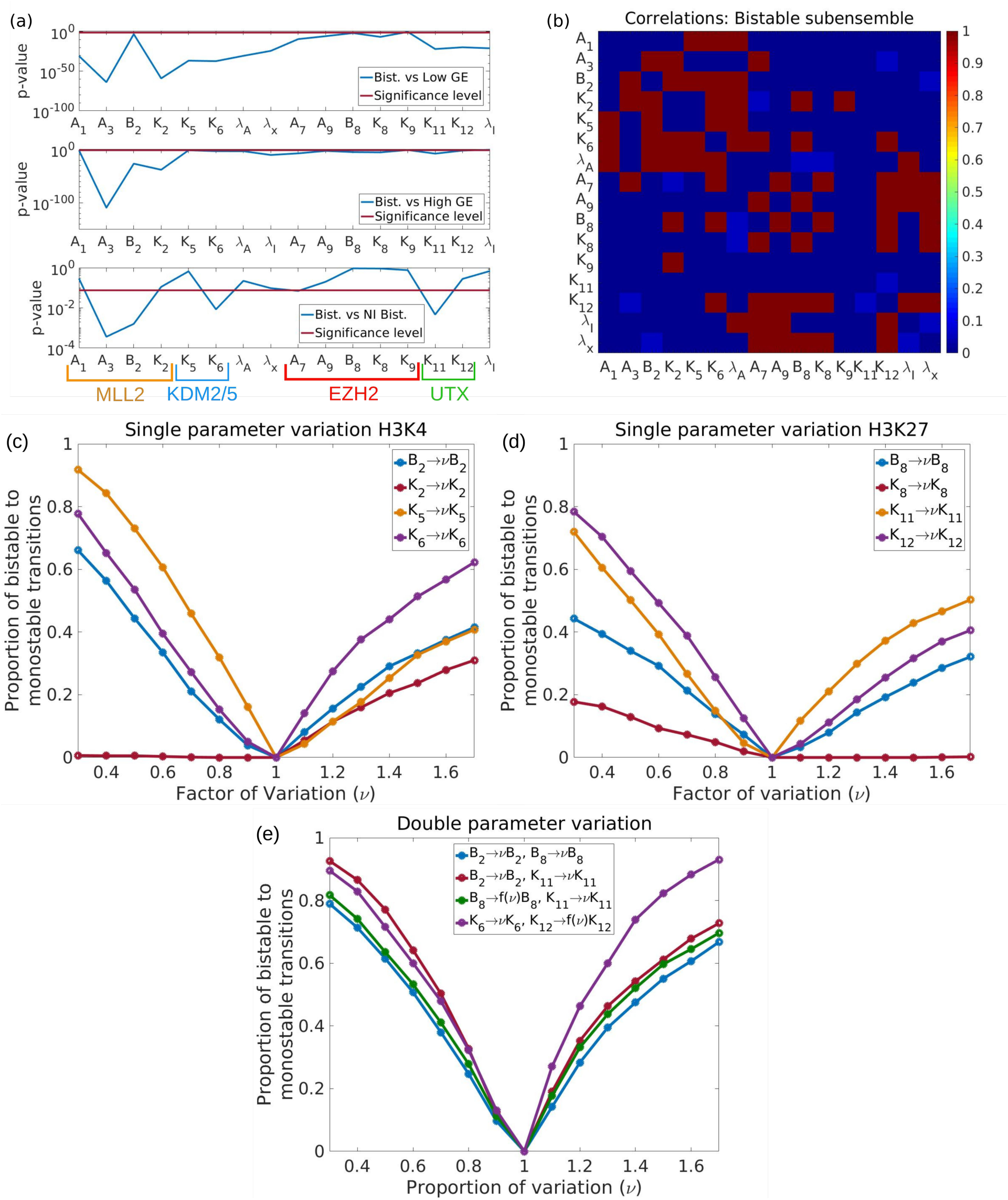
Tackling heterogeneity via parameter sensitivity analysis. Plot (a): *p*-values of the Kolmogorov-Smirnov comparison of the marginal distribution for each component of *θ* = (*A*_1_, *A*_3_, *B*_2_, *K*_2_, *K*_5_, *K*_6_, *λ*_*A*_, *λ*_*x*_, *A*_7_, *A*_9_, *B*_8_, *K*_8_, *K*_9_, *K*_11_, *K*_12_, *λ*_*I*_) corresponding to each of the sub-ensembles (see S1 File, Section S.1.4.2 for details). Plot (b): correlations with the Bistable sub-ensemble. Red squares correspond to pairs such that the null hypothesis that the parameters are uncorrelated can be rejected (p-value¡0.05), dark blue, corresponds to p-values¿0.5 (uncorrelated pairs), and light blue, to p-values≳0.05, i.e. pairs of parameters for which the evidence of absence of correlations is weaker. Plots (c) to (e) show the sensitivity analysis corresponding to the Bistable sub-ensemble. Recall that, according to our validation results, we are identifying the systems within this sub-ensemble with ESCs or undifferentiated cell states. Plots (c), (d), and (e) show the sensitivity to changes in kinetic parameters affecting the enzymes modifying H3K4, H3K27, and several combinations, respectively.

We characterise heterogeneity within the (sub)ensemble using two statistics, namely, the posterior marginal PDF for each of the parameters (see Figure S.3 in S1 File) and the linear correlations between them. The former allows us to identify systematic, statistical significant bias in the values of each of the parameters associated with an specific behaviour (in this case, bistability) [33, 34]. In Figure 6(a), we plot the *p* value corresponding to the Kolmogorov-Smirnov (KS) test comparing the marginal PDFs of the bistable sub-ensemble with those of the Low GE, High GE and noise-induced bimodal sub-ensembles. Provided that *p* is smaller than the significance level, the smaller the *p*-value, the more statistically significant is the difference between the marginal PDFs. Such information allows us to identify the subset of parameters which are more likely to drive the behaviour of the system from one sub-ensemble (e.g. bistable) into another (e.g. Low GE or High GE).

In order to perform our analysis, we choose the subset of parameters for which the KS test gives smaller *p*-values for the comparison of the bistable sub-ensemble with the Low GE and the High GE sub-ensembles: *A*_3_, *B*_2_, *K*_2_, *B*_8_, *K*_8_, *K*_11_, and *K*_12_ (see Figure 6(a)). The motivation for such a choice is that this set of parameters are the ones that are more relevant to bistable behaviour [33, 34]. We have analysed the response of the bistable sub-ensemble by measuring the proportion of bistable *θ*s that become monostable when these parameters are modified (one at a time) by a factor *ν* (see Figure 6(d)). According to such a metric, our results show that the parameters that fare better concerning their ability to beat heterogeneity are the rescaled Michaelis-Menten parameters for the KDMs (i.e. *K*_5_ and *K*_6_ for KDM2/KDM5 which remove me3 from trimethylated H3K4 residues and *K*_11_ and *K*_12_ for KDM6 which remove trimethyl from H3K27me3). By contrast, our analysis shows that targeting the activity of MLL2 or EZH2 is a much poorer strategy regarding the aim of producing a maximally homogeneous sub-ensemble response.

Single parameter variations have limitations of two kinds. First, when considering *ν* > 1, the response curve is nonlinear but very quickly saturates to a response between 35%-65%, approximately (see Figures 6(c) and (d)). By contrast, if *ν* < 1, the response curve is approximately linear. Whilst this scenario is more favourable than the *ν* > 1 one, ideally we would like to have a combination of both: a sigmoidal response curve with saturation value close to 100%. Such scenario is achieved when considering combinations of parameters (see Figure 6(e)). Specifically, of the combinations that we have explored, the most efficient combination is *K*_6_-*K*_12_, i.e. those involving both parameters regulating KDM activity. The reason for the enhanced efficiency of this specific combination is that *K*_6_ and *K*_12_ are correlated (see Figure 6(b)). All other combinations that we have tried involved uncorrelated parameters and perform worse than the *K*_6_-*K*_12_ one.

## Conclusion

Tumor cell heterogeneity is a major barrier for efficient design of targeted anti-cancer therapies. A diverse distribution of phenotypically distinct tumour-cell subpopulations prior to drug treatment predisposes to non-uniform responses, leading to the elimination of sensitive cancer cells whilst leaving resistant subpopulations unharmed. Despite tumour cell heterogeneity has been recognized as a *bona fide* engine for drug resistance [1, 3, 76], few successful approaches [17] have been proposed aimed at formulating strategies capable of quantifying the variability associated to individual cancer cell heterogeneity and minimizing its undesirable impact on clinical outcomes. Our current work accepts this challenge to provide a computational approach that allows the rational design of combinatorial therapies involving epigenetic drugs against cancer-driving chromatin modifiers [77–79].

To analyse the effects of targeted therapies which affect the enzymatic efficiency of specific chromatin modifiers and combinations, thereupon we have formulated a stochastic model of a bivalent transcription factor [42, 43, 45, 58], for which we have been able to derive the steady-state probability distribution function *via* singular perturbation and model reduction analysis. This result allows us to characterise the different qualitative behaviours of the system (open (High GE), closed (Low GE), bistable, and noise-induced bistability). In order to tackle heterogeneity, we follow [33, 34] and generate an ensemble of models which we then classify into four sub-ensembles according to their qualitative behaviour. Comparison between analytical results and experimental data allows us to determined that the so-called Bistable and the High GE sub-ensembles show the same behaviour as undifferentiated and differentiated cell types, respectively. By contrast, the Low GE sub-ensemble failed to show any behaviour that could be identified as any cell type previously studied in [20, 21].

Since undifferentiated cells, i.e. locked in a self-renewing phenotype, have been identified as aberrant phenotypes which, if left unchecked, will eventually give rise to a tumour [21, 25, 33, 34], we have focused on analysing the role of heterogeneity within the Bistable sub-ensemble regarding their response to targeted epigenetic therapies. Such therapies are assumed to affect the value of specific parameters which, in turn, alter the enzymatic activity of the corresponding chromatin modifiers. Although our ensemble approach provided a rationale to choose those parameters to which undifferentiated (bistable) behaviour should be more sensitive [33, 34], it was not surprising that single-targeted strategies mostly failed to circumvent the therapeutic problems represented by tumour heterogeneity whereas combinatorial strategies fared much better. Specifically, those strategies involving the H3K4 and H3K27 methyltransferases MLL2 and EZH2 were predicted to be notably less effective than those involving modulators of the histone H3K4 and H3K27 demethylases KDM5 and KDM6A/UTX. Accordingly, we are accumulating evidence that transcriptional differentiation programs governed by KDM6A/UTX can enforce and safeguard cellular identity in several cancer types whereas its pharmacological manipulation might impede tumour progression to highly aggressive subtypes by blocking epigenetic roads to de-differentiation [80–82].

Our theoretical framework provides a coherent basis for the development of an *in silico* platform capable of identifying the epigenetic drugs combinations best-suited to therapeutically manage non-uniform responses of heterogenous cancer cell populations. The proposed paradigm might accelerate future tool developments on epi-drugs applications with capability for patient stratification based on the dominant cell subtype in individual tumours.

## Supporting information

### S1 File. Supplementary material

The file contains a more detailed description of the model formulation and model analysis, including a full derivation of the asymptotic model reduction.

## Acknowledgments

T.A., J.A.M., and J.S. have been partially funded by the CERCA Programme of the Generalitat de Catalunya. The authors also acknowledge support from State Research Agency (grants MTM2015-71509-C2-1-R, MTM2015-71509-C2-2-R, RTI2018-098322-B-I00, PGC2018-098676-B-I00 and SAF2016-80639-P) and AGAUR (projects 2014SGR229, 2017SGR1049 and 2017SGR01735). J.S. has been also funded by a “Ramón y Cajal” Fellowship (RYC-2017-22243).

